# Single-cell transcriptional analysis of the immune tumour microenvironment during myeloma disease evolution

**DOI:** 10.1101/2021.10.22.464971

**Authors:** Danielle C. Croucher, Laura M. Richards, Daniel Waller, Zhihua Li, Xian Fang Huang, Marta Chesi, P. Leif Bergsagel, Michael Sebag, Trevor J. Pugh, Suzanne Trudel

## Abstract

Multiple myeloma is universally preceded by a premalignant disease state. However, efforts to develop preventative therapeutic strategies are hindered by an incomplete understanding of the immune mechanisms associated with progression. Using single-cell RNA-sequencing, we profiled 104,880 cells derived from the bone marrow of Vκ*MYC mice across the myeloma progression spectrum, of which 97,720 were identified as non-malignant cells of the tumour microenvironment. Analysis of the non-malignant cells comprising the immune microenvironment identified mechanisms associated with disease progression in innate and adaptive immune cell populations. This included activation of IL-17 signaling in myeloid cells from precursor mice, accompanied by upregulation of Il6 gene expression in basophils. In the T/Natural killer cell compartment, we identified Tox-expressing CD8+ T cells enriched in the tumour microenvironment of mice with overt disease, with co-expression of LAG3 and PD-1, as well as elevated T cell exhaustion signatures in mice with early disease. We subsequently showed that early intervention with combinatorial blockade of LAG3 and PD-1 using neutralizing monoclonal antibodies delayed tumor progression and improved survival of Vκ*MYC mice. Together, this work provides insight into the biology of myeloma evolution and nominates a treatment strategy for early disease.

## Introduction

Multiple myeloma (MM) is a hematological malignancy characterized by accumulation of clonal, terminally-differentiated plasma cells in bone marrow (BM). Previous studies have confirmed that MM is universally preceded by premalignant precursor disease states [1, 2], including monoclonal gammopathy of undetermined significance (MGUS) and smoldering MM (SMM). However, there are no reliable methods to predict if or when precursor patients will progress to overt disease. Therefore, a better understanding of the molecular processes contributing to progression is needed to improve surveillance, clinical management and treatment.

The challenge of risk-stratifying patients with precursor disease has been approached using several different methods. Kauffmann et al. used serial BM specimens to show that primary driver events such as the recurring immunoglobulin translocations are not sufficient to drive progression, as these alternations are present in both MGUS with stable disease and MM patients [3]. Similarly, while Lopez-Corral et al. detected significantly more copy number variants (CNVs) in MM compared to MGUS/SMM, alternations that appeared exclusive to MM, such as 1q+, 21q+, del16q, and del22q, were subsequently shown to be present in plasma cell subclones at the precursor stages [4]. These data suggest that most, if not all chromosomal aberrations are already present in the early stages of disease. Similarly, exome analysis of paired precursor and MM samples demonstrated that most somatic mutations in MM predate the onset of clinical malignancy [5], as does intratumoural heterogeneity within the malignant cell compartment [6]. Thus, there is significant evidence that somatic mutations and CNVs alone cannot account for MM disease evolution.

Gene expression profiling is an alternative readout to explore the molecular underpinnings of high-risk precursor disease. Transcriptional signatures derived from bulk BM plasma cell samples have been generated to define high-risk MM [7], as well as to risk-stratify patients with precursor disease [8]. More recently, gene expression profiling at single-cell resolution has been used as a tool to precisely define the transcriptional heterogeneity of precursor and myeloma tumours. These studies have revealed substantial intratumoural heterogeneity and defined subclonal transcriptional populations [9], highlighting the value of exploring gene expression at the single-cell level. However, the fact remains that most tumours from MGUS and SMM patients exhibit clinical stability in spite of advanced genomic complexity and subclonal evolution. Therefore, progression to overt MM may also depend on factors that are extrinsic to tumour cells, including the immune cells that comprise the tumour microenvironment (TME). This is supported by studies demonstrating progressive growth of tumour cells derived from clinically-stable precursor patients upon xenotransplantation into a humanized mouse model, thereby bypassing host immunosurveillance [10]. More recently, a key role for the cytokine IL-17 has been identified in promoting MM progression, with high IL-17 levels predictive of progression from SMM to malignant MM [11].

In this study, we sought to understand how immune cell populations in myeloma change throughout disease evolution using single-cell RNA-sequencing (scRNA-seq) and, to then nominate novel mechanisms that drive tumour progression. To enable study of a longitudinal, genetically-uniform cohort of subjects, we employed a genetically-engineered, immune-competent mouse model of MM (Vκ*MYC), which is biologically recapitulative of MM disease progression through precursor stages [12]. Vκ*MYC mice develop a progressive accumulation of clonal BM plasma cells that, as in the human disease, can be monitored indirectly by measuring monoclonal serum immunoglobulins (M-protein) in blood. Herein, we present a comprehensive single-cell analysis of the natural progression of the immune TME in myeloma using this high fidelity model of human disease.

## Results

### Composition of the immune TME throughout disease progression in Vκ*MYC mice

To characterize how the immune TME changes during disease progression in Vκ*MYC mice, we performed scRNA-seq profiling of unselected BM cells from mice at various stages of progression (Figure 1A). After removal of low-quality cells and predicted doublets, our data set contained 104,880 single cell expression profiles including 85,269 cells from a disease stage-spanning cohort of Vκ*MYC mice (n=15) and 19,611 cells from age-matched C57BL/KaLwRij control mice (n=3). Unsupervised clustering of all 104,880 cells revealed 42 transcriptionally distinct clusters (Figure 1B).

**Figure 1.**
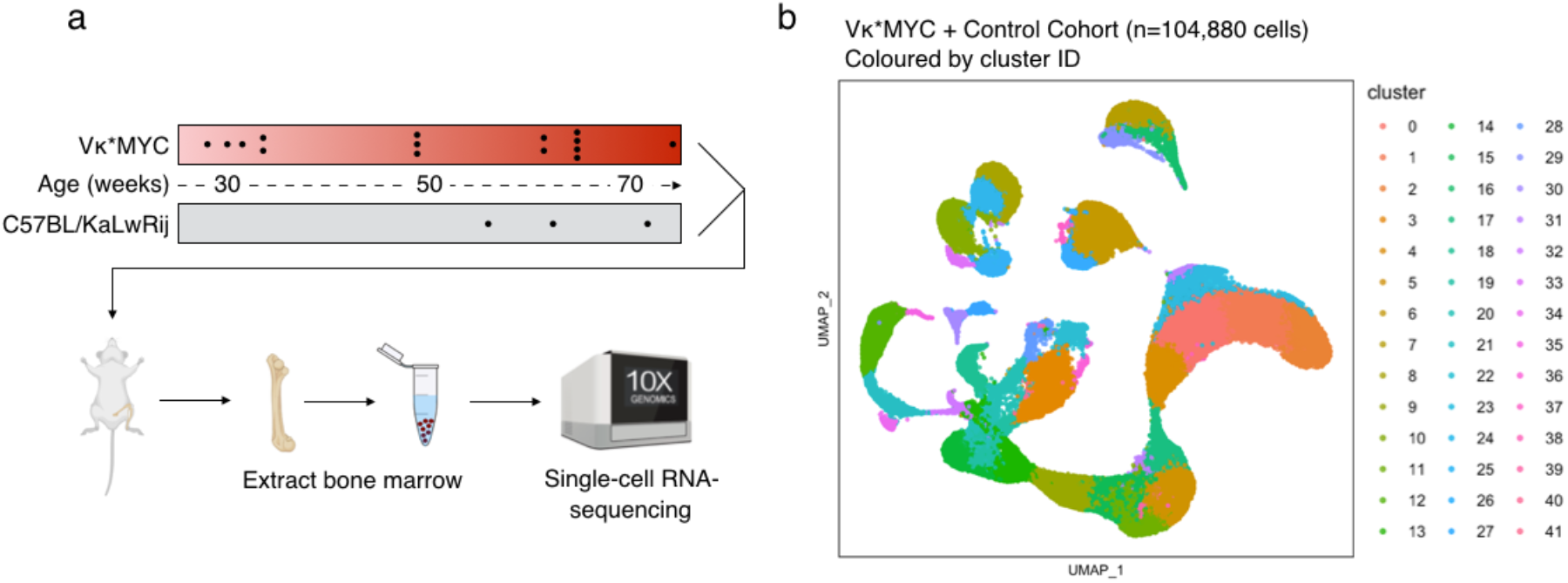
Single-cell profiling of the immune TME landscape in Vκ*MYC mice. (a) Overview of study design, cohort composition, sample workflow and analysis strategy. (b) UMAP visualization of 104,880 cells from BM of full study cohort, coloured by transcriptionally-defined population.

We then assigned these clusters to major cell types using expression of known marker genes and reference-driven annotations from SingleR [13] (Figure 2A-C). As expected, we detected various non-immune populations resident to the BM such as erythroid and progenitor cells (C9, C12, C13, C15, C21, C32), as well as malignant plasma cell populations (C7, C16, C29). However, given the role of immune dysregulation in myeloma pathogenesis, we focused our analysis on immune cell populations, which included T/Natural killer (NK) cells (C6, C26, C37), B lymphocytes (C8, C10, C24, C25, C33), monocytes/macrophages (C3, C14, C19, C20, C22, C28, C34, C35, C36, C40, C41), plasmacytoid dendritic cells (pDC) (C18), neutrophils (C0, C1, C2, C4, C5, C11, C17, C23, C31, C38, C39), and basophils (C27, C30) (Figure 2B). Thus, our analysis of the immune TME during progression encompassed single-cell transcriptional profiles for 84,009 immune cells.

**Figure 2.**
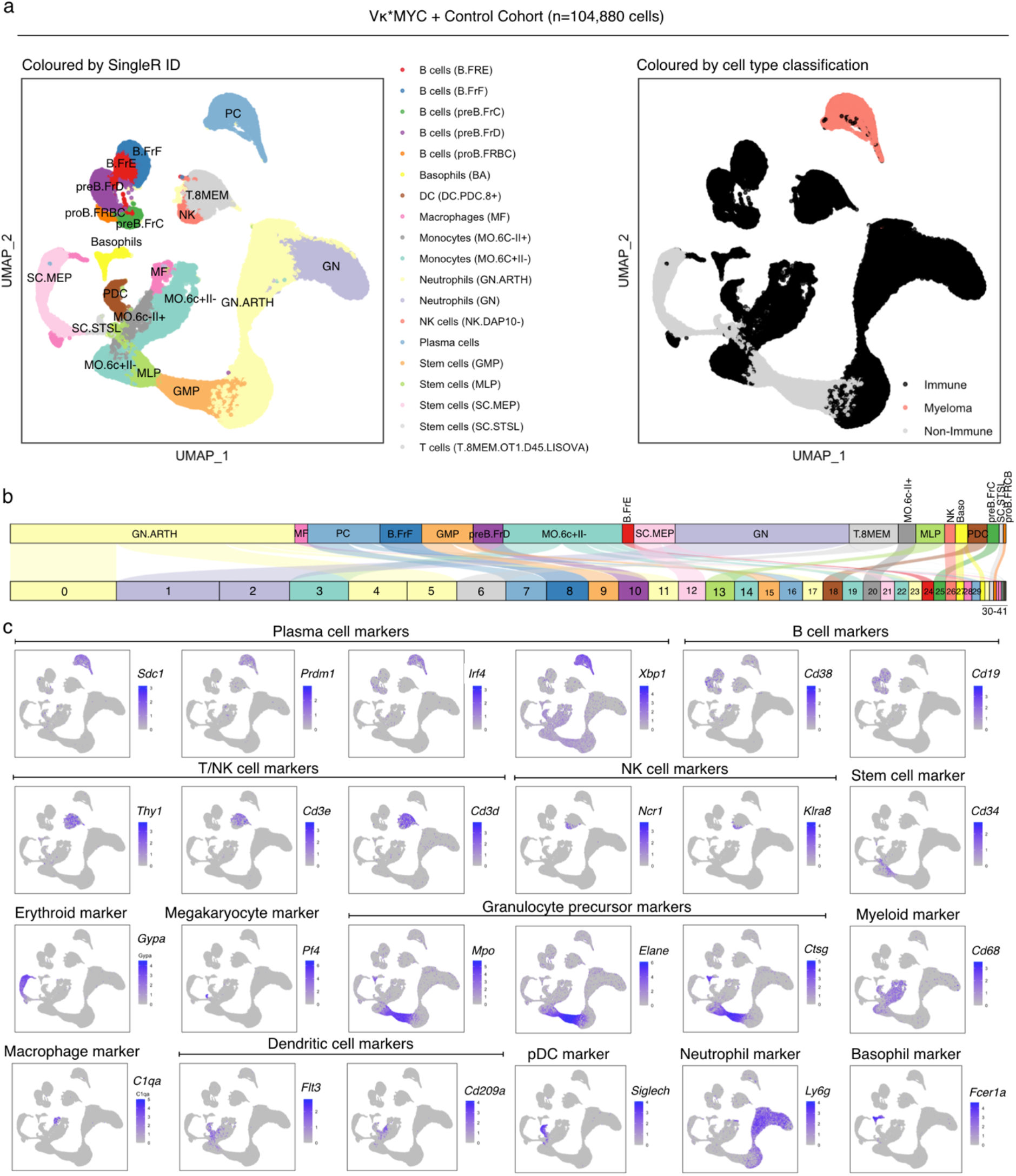
Cellular composition of the immune TME. (a) UMAP visualization of 104,880 cells from BM of full study cohort, coloured by lineage annotation (left), and cell type classification (right). (b) Flow diagram depicting assignment of clusters (bottom track) to lineage annotation (top track). Colours correspond to SingleR ID from (a) and are scaled to the number of cells within each cell type/cluster. (c) UMAP visualization of 104,880 cells from BM of full study cohort, coloured by relative expression of depicted marker genes.

To explore the relationship between immune cell composition and disease progression, we calculated the correlation between the relative proportion of each immune cell cluster within its respective lineage and disease burden as measured by % malignant cells in the BM. As shown in Figure 3A, our analysis revealed that disease progression is associated with a decrease in pDCs (Figure 3B; C18, Pearson’s r=−0.566, P=0.028) and an increase in the C28 population of macrophages (Figure 3C; Pearson’s r=0.682, P=0.006). Although the granulocyte cluster, C38, was also associated with disease progression (Pearson’s r=0.602, P=0.018), closer inspection of the data revealed that enrichment was driven by only one Vκ*MYC mouse (Figure 3D) and that re-computation without this sample rendered the correlation not significant (Pearson’s r=0.359, P=0.207). Nonetheless, the observed compositional shifts in pDCs during progression in our analysis is consistent with the recently reported scRNA-seq study of MGUS, SMM and MM patients by Zavidij et al. [14] and thus, supports the validity of Vκ*MYC mice to recapitulate myeloma precursor biology. Moreover, it implicates myeloid cell populations in particular, with disease progression, providing rationale for a more in depth analysis of this cell lineage.

**Figure 3:**
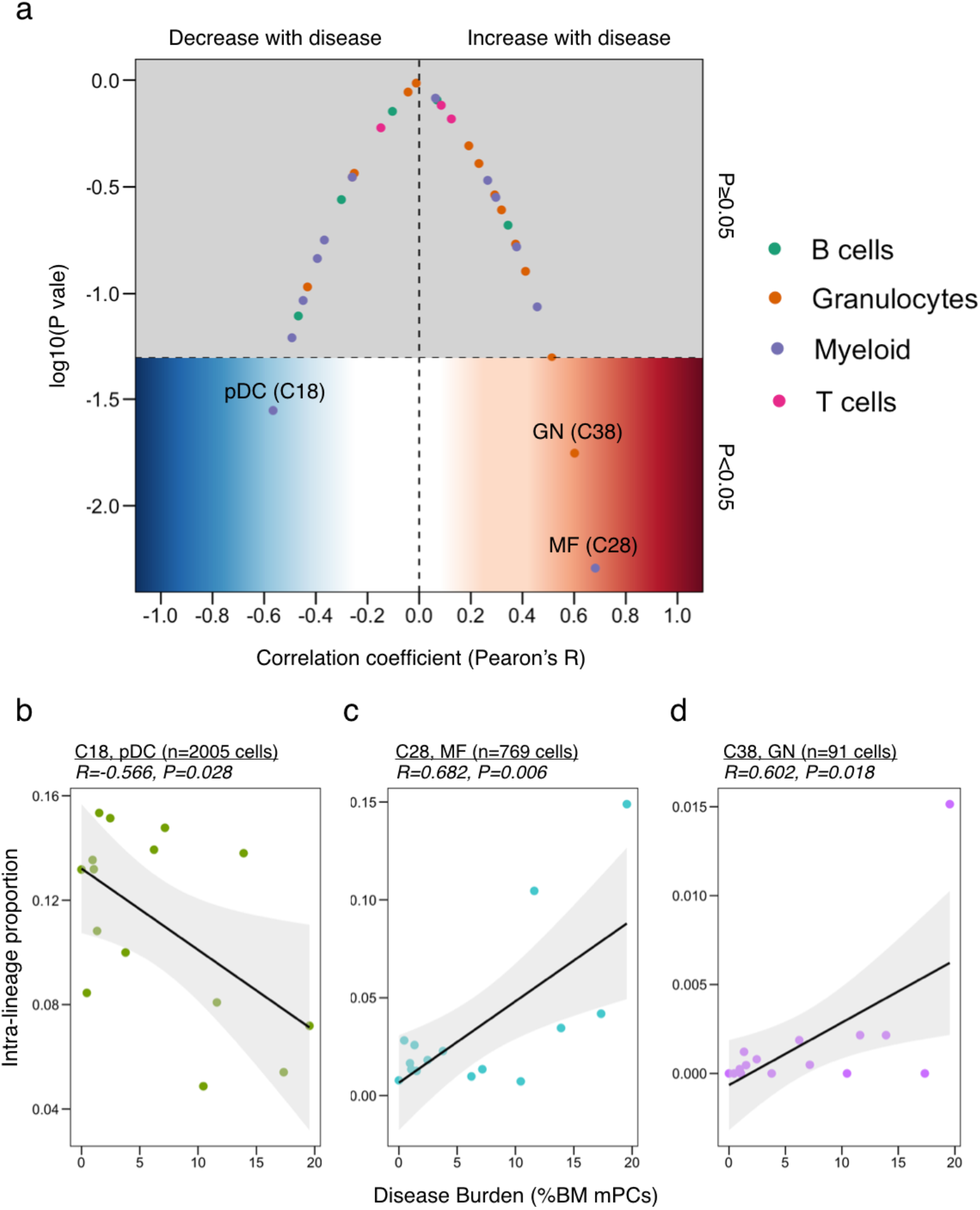
Compositional changes in the immune TME throughout progression. (a) Correlation plot relating disease burden to immune TME composition using the full set of scRNA-seq profiles. For each sample, the number of cells in each cluster was divided by the total number of cells in that lineage. Each point reflects a cluster-specific test for association (Pearson’s R) between the calculated lineage-wise proportion and the total number of malignant plasma cells (mPCs) detected from scRNA-seq analysis, coloured by lineage. (b-d) Scatter plots revealing sample-level relationship between disease burden and immune TME composition for clusters with significant correlation from (a) including (b) C18, (c) C28, and (d) C38. Each data point represents one sample.

### Sub-clustering analysis of the myeloid compartment implicates IL-17 signaling in the intermediate stages of myeloma progression

To investigate myeloid cell subtypes and states in more detail, we re-clustered a subset of our single-cell expression data corresponding to cells of the myeloid lineage (n=13,594 cells) (Figure 4A). The resulting 23 myeloid sub-clusters were then annotated based on expression of known myeloid marker genes including Ly6c-monocytes (*Itgax*+, *S1pr5*+, *Spn*^HI^, *Sell*^HI^), Ly6c+ monocytes (*Ccr2*+, *Spn*^LO^, *Sell*^LO^), pDCs (*Siglech*+), classical DCs (*Flt3*+, *Cd209*+), macrophages (*C1qa*+), interferon-gamma-response monocytes (*Cxcl10*+, *Ifit1*+, *Ifit2*+, *Ifit3*+) and cell cycle high monocytes (*Mki67*^HI^) (Figure 4B).

**Figure 4:**
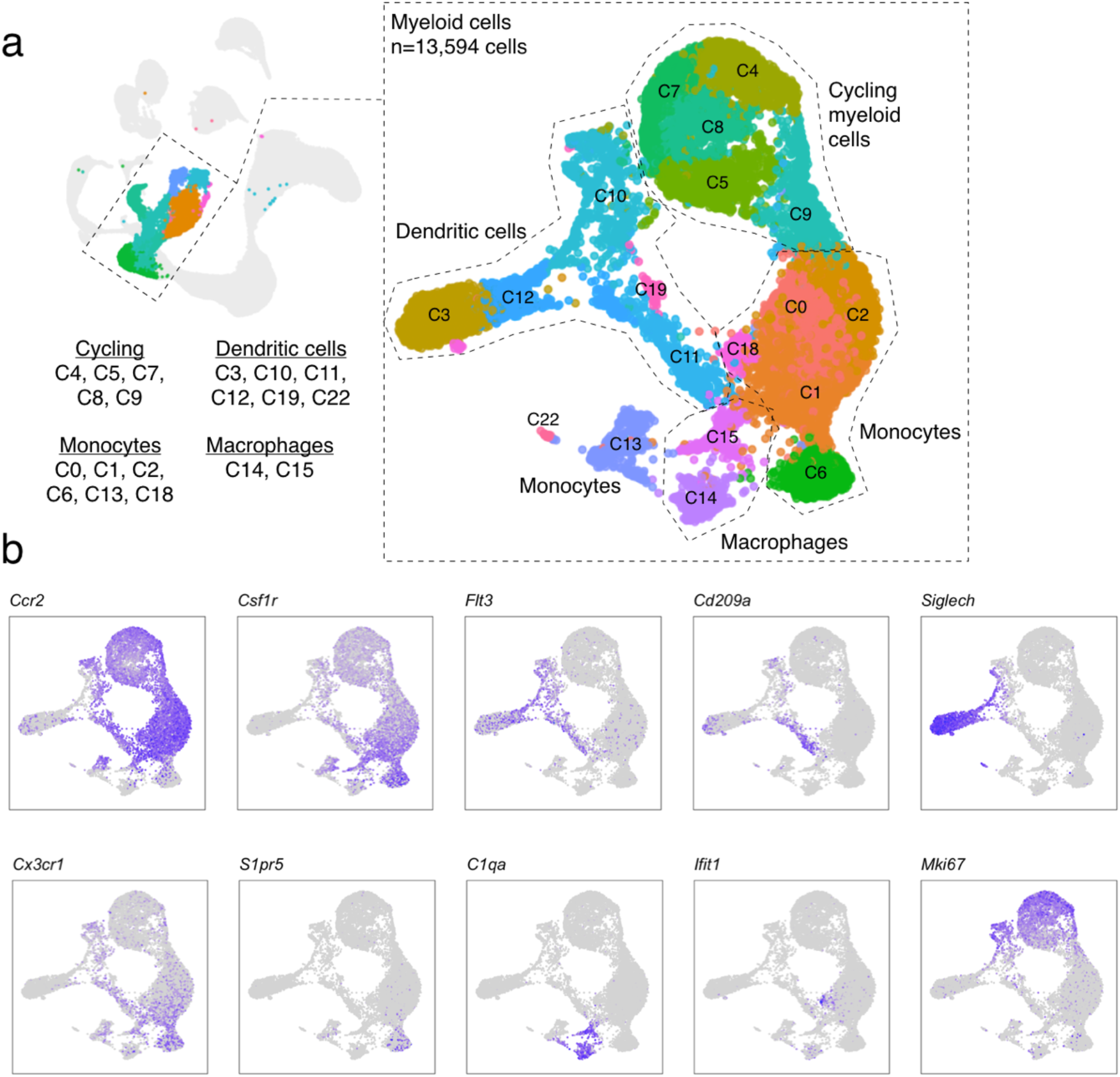
Transcriptional heterogeneity within the myeloid lineage as determined by scRNA-seq. (a) Subclustering analysis of 13,594 myeloid cells revealing transcriptionally-distinct populations as visualized using UMAP plot. Each population is labeled by a specific colour/number and annotated using (b) select myeloid marker genes.

To determine whether any of these clusters were enriched at a particular stage of disease, we assigned mice to one of three disease stage groups based on age and serum M-protein levels (Figure 5A-B), which ultimately included 5 Vκ*MYC mice without detectable disease (early-MM, EMM1-5, 27-31 weeks), 3 Vκ*MYC mice with intermediate disease (int-MM, IMM1-3, 49 weeks), and 3 Vκ*MYC mice with active MM (active-MM, AMM1-3, 61-74 weeks), plus the 3 age-matched control C57BL/KaLwRij mice (Control, Cont1-3, 55-72 weeks). The C14 macrophage cluster, which corresponds to C28 from Figure 1B and Figure 2B, was almost exclusively found in mice with active-MM, consistent with the association of these cells with disease burden (Figure 5C). Moreover, the expression profile of these cells is consistent with that of CD16+ monocytes described by Zavidij et al. [14] including upregulation of *Ms4a7*, *Lst1*, *Aif1*, *Fcer1g* and *Fcgr4* (Figure 5D). Thus, our data provide additional support for the association of these cells with disease progression and provide further rationale for future work to explore their role in myeloma biology.

**Figure 5:**
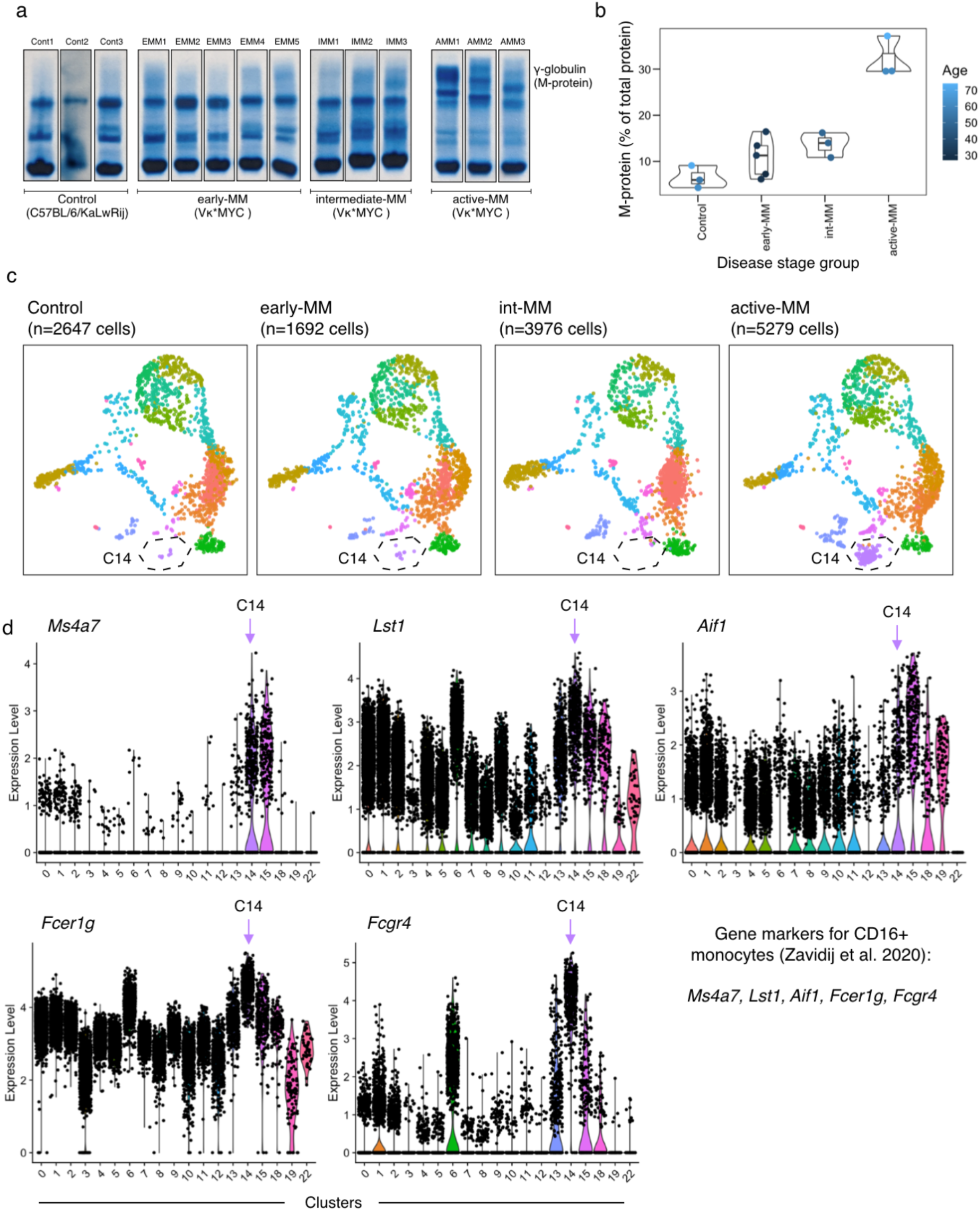
Defining myeloid subpopulations associated with defined disease stage groups. (a) SPEP gels depicting M-protein bands (γ-globulin) across defined disease stage groups. (b) Disease burden across disease stage groups as determined by quantification of M-protein measurements from SPEP. Each point represents one mouse and is coloured by age of mouse at the time of sacrifice. (c) Disease stage-specific UMAP plots from myeloid subclustering analysis, highlighting cluster containing putative CD16+ monocytes (C14). (d) Violin plot depicting expression of CD16+ monocyte marker genes from Zavidij et al. [81] across myeloid clusters from Vκ*MYC mouse cohort.

Our focused subclustering analysis of myeloid subtypes and cell states also revealed a significantly different distribution of Ly6c+ monocyte clusters amongst Vκ*MYC mice (Figure 6A-B), with enrichment of C0 in int-MM compared to early-MM and active-MM (Figure 6B, p=0.026 and p=0.003, respectively). Therefore, Ly6c+ monocytes from mice with int-MM demonstrate a distinct transcriptional shift during progression. To further investigate the heterogeneity of Ly6C+ monocyte cellular states and the association between these programs and disease progression, we explored the expression profiles of all Ly6c+ monocyte clusters (C0, C1, C2). Using differential expression analysis, we found 31, 52 and 12 marker genes upregulated in C0, C1 and C2, respectively (Figure 6C), and then investigated the pathways associated with the 31 genes significantly upregulated in int-MM-associated C0 cells with gene set enrichment analysis. Accordingly, one of the more highly enriched pathways in C0 was related to IL-17 signaling (Figure 6D, FDR=0.000643), which is of interest given that higher IL-17 cytokine in the BM is associated with acceleration of disease progression in SMM patients [158].

**Figure 6:**
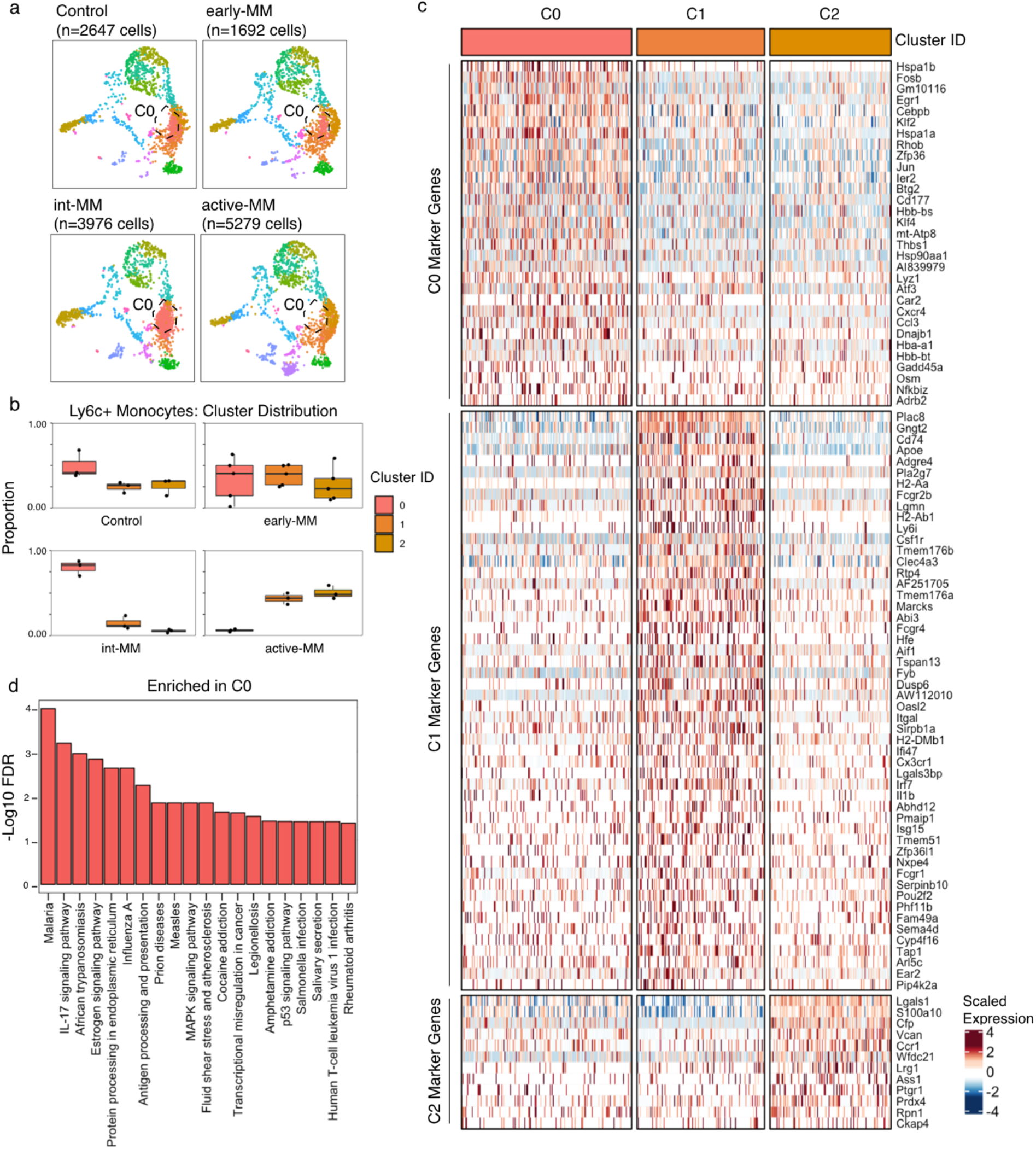
Precursor disease is marked by elevated IL-17 signaling in the immune TME. (a) Disease stage-specific UMAP plots from myeloid subclustering analysis, highlighting Ly6c+ monocyte cluster differentially enriched throughout progression (C0). (b) Ly6c+ monocyte cluster distribution across control, early-MM, int-MM and active-MM disease stages. Each point represents the proportion of the indicated cluster within all Ly6c+ monocytes for a given sample. (c) Heatmap depicting marker genes for each of three Ly6c+ monocyte clusters as determined by differential expression analysis. Data represent scaled expression values (any values outside a range of −2.5 to 2.5 were clipped). (d) Top 20 positively enriched terms from Enrichr gene set enrichment analysis (FDR<0.05) computed using C0 marker genes from (c).

Mechanistically, it has been shown that IL-17 in this context induces IL-6 expression in eosinophils [11], which is a cytokine with well-established, pro-survival effects on myeloma cells [15, 16]. Assuming that activation of IL-17 signaling in C0 cells reflects the presence of IL-17 in the TME of Vκ*MYC mice, we explored the possibility of IL17-induced IL-6 expression in C0 cells, but observed no *Il6* gene expression in any myeloid cells from our data set (Figure 7A). However, analysis of all 104,880 TME cells from our data set revealed that *Il6* was highly expressed in basophils (Figure 7B-C), which, like eosinophils, belong to the polymorphonuclear leukocyte lineage. Moreover, *Il6* expression in basophils positively correlated with the same IL-17 signaling gene signature identified in C0 (Figure 7D, R=0.504, P<2.2e-16). Thus, our data reveal that a link between IL-17 signaling and *Il6* expression, previously only reported in eosinophils [11], is also active in basophils. Furthermore, it demonstrates that this IL-17-basophil-IL-6 transcriptional axis is specifically relevant during the int-MM transition phase of disease.

**Figure 7:**
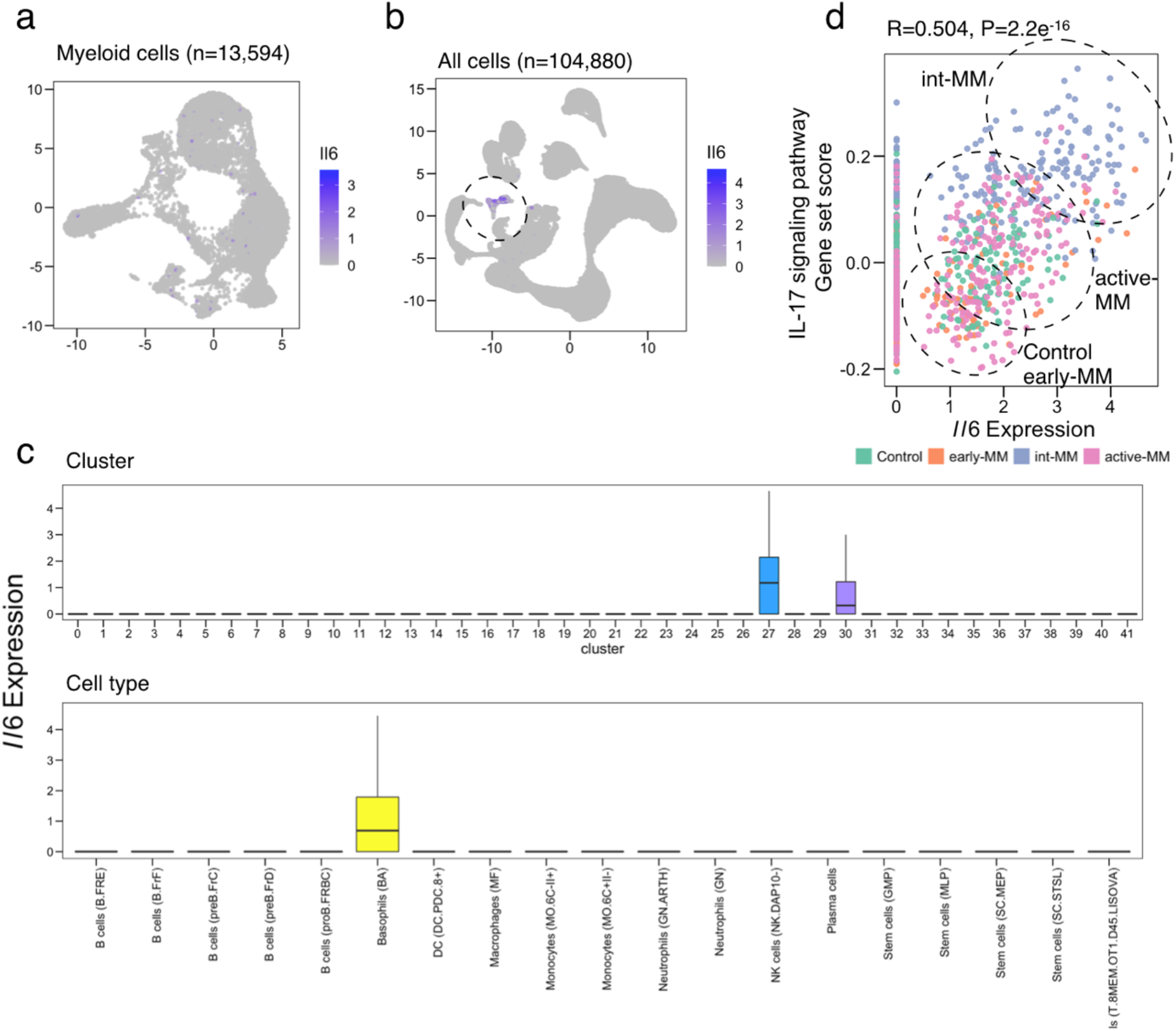
Single-cell analysis establishes novel IL17-basophil-IL6 transcriptional axis in the precursor immune TME. (a) UMAP visualization of log-normalized *Il6* expression in myeloid cells. (b) UMAP visualization of log-normalized *Il6* expression in all BM cells. (c) Box plots depicting distribution of *Il6* expression in full BM data set, grouped by transcriptional cluster (top) and cell type/lineage annotation (bottom). (d) Relationship (Pearson’s R) between IL-17 signaling gene set score and *Il6* expression in basophils (n=958), coloured by disease stage group.

### T cell exhaustion occurs early in Vκ*MYC mice and is accompanied by expression of multiple immune checkpoint receptors

Given the direct role for T cells in mediating anti-tumour effector functions, we next performed an in-depth re-clustering analysis of the T/NK cell compartment using the same cohort of mice described above (n=5,101 cells). This analysis yielded 15 subpopulations (Figure 8A), which we annotated as the following cell subtypes using a combination of known marker genes [17, 18] (Figure 8B): Naïve T cells (*Cd3*+, *Ccr7*+, *Sell*+), CD4 Helper T cells (*Cd3*+, *Cd4*+), Regulatory T (Treg) cells (*Cd3*+, *Cd4*+, *Foxp3*+), T Helper 17 (Th17: *Cd3*+, *Il17a*+), CD8 T cells (*Cd3*+, *Cd8a*+), NK cells (*Cd3d*−, *Cd3e*−, *Ncr1*+), Innate Lymphoid Cells (ILC), Type 2 (ILC2: *Cd3*-, *Rnf128*+), and ILC3 (*Cd3*−, *Ncr1*+, *Klrb1b*+).

**Figure 8:**
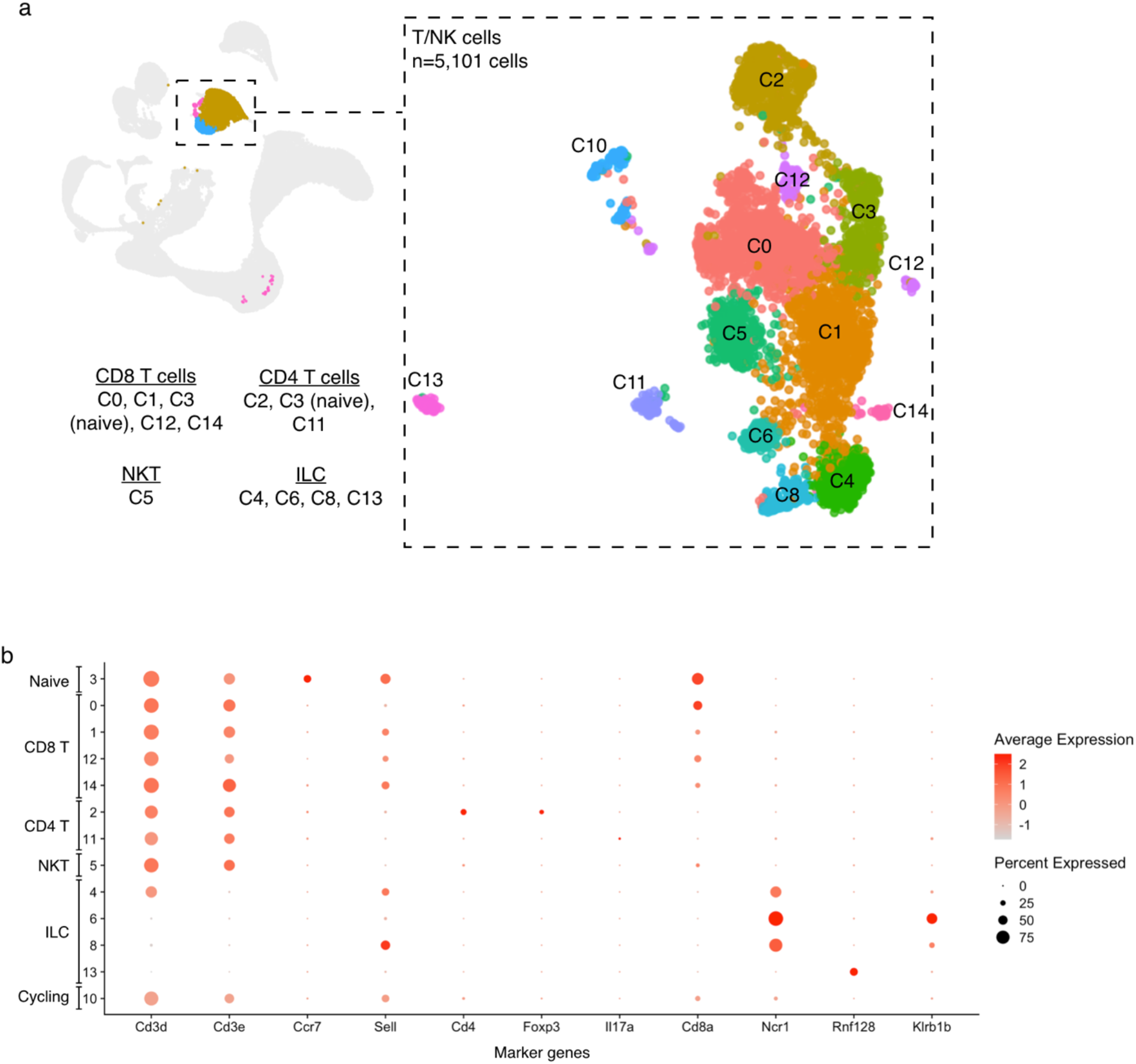
Unsupervised subclustering of 5,101 T/NK cells reveals cell subtypes from scRNA-seq data. (a) Subclustering analysis of 5,101 T/NK cells revealing transcriptionally-distinct populations as visualized using UMAP plot. Each population is labeled by a specific colour/number and annotated using (b) select T/NK marker genes. Dot plot in (b) depicts expression of T cell-related marker genes (y-axis) across clusters (x-axis), which are ordered/grouped by similar cell subtypes for ease of interpretation. The size of the dot encodes the percentage of cells within a cluster, while the color encodes the average expression level across all cells within a cluster.

To determine whether T/NK cell subtypes or transcriptional programs are associated with disease progression, we evaluated the proportion of each cluster across disease stages. The CD8 T cell cluster, C0, was significantly enriched for cells from mice with active-MM compared to control (P=0.0154), early-MM (P=0.0147) and int-MM groups (P=0.0171) (Figure 9A). To identify marker genes that define the transcriptional program of these cells, we performed differential expression analysis between C0 and all other T/NK clusters, which revealed only 12 genes to be significantly upregulated in C0 T cells (Figure 9B). Interestingly, this included two transcriptional factors associated with T cell exhaustion, *Tox [19]* and *Eomes [20]*, suggesting that the immune TME of mice with active-MM is enriched for T cells with an exhausted phenotype. Given the role of immune checkpoint receptors (ICRs) in mediating T cell exhaustion [21], we explored the expression of various ICRs on C0 cells including *Btla* (CD272), *Cd244*, *Ctla4*, *Entpd1* (CD39), *Havcr2* (TIM-3), *Lag3*, *Pdcd1* (PD-1) and *Tigit*. This analysis revealed that the most highly enriched ICRs on C0 cells were *Lag3* (8.86%) and *Pdcd1* (8.13%), suggesting that exhaustion may be mediated by co-expression of these ICRs (Figure 9C). Interestingly, *Lag3* and *Pdcd1* expression was enriched in cells from both int-MM and active-MM (Figure 9D) suggesting that co-expression of LAG3 and PD-1 cell surface receptors occurs earlier in disease progression. Flow cytometric analysis confirmed a strong positive association between disease progression and LAG3/PD-1 co-expression on CD8+ T cells in the Vκ12598 transplantable model of Vκ*MYC (Figure 10A, R = 0.950, P=4.484e-11). Moreover, using our scRNA-seq data, we found that CD8+ T cells from mice with int-MM and active-MM displayed high levels of exhaustion (Figure 10B), as did cells co-expressing *Pdcd1* and *Lag3* (Figure 10C-D). Thus, these results support that T cell exhaustion occurs early in the course of disease progression and may be mediated by multiple ICRs during disease evolution.

**Figure 9:**
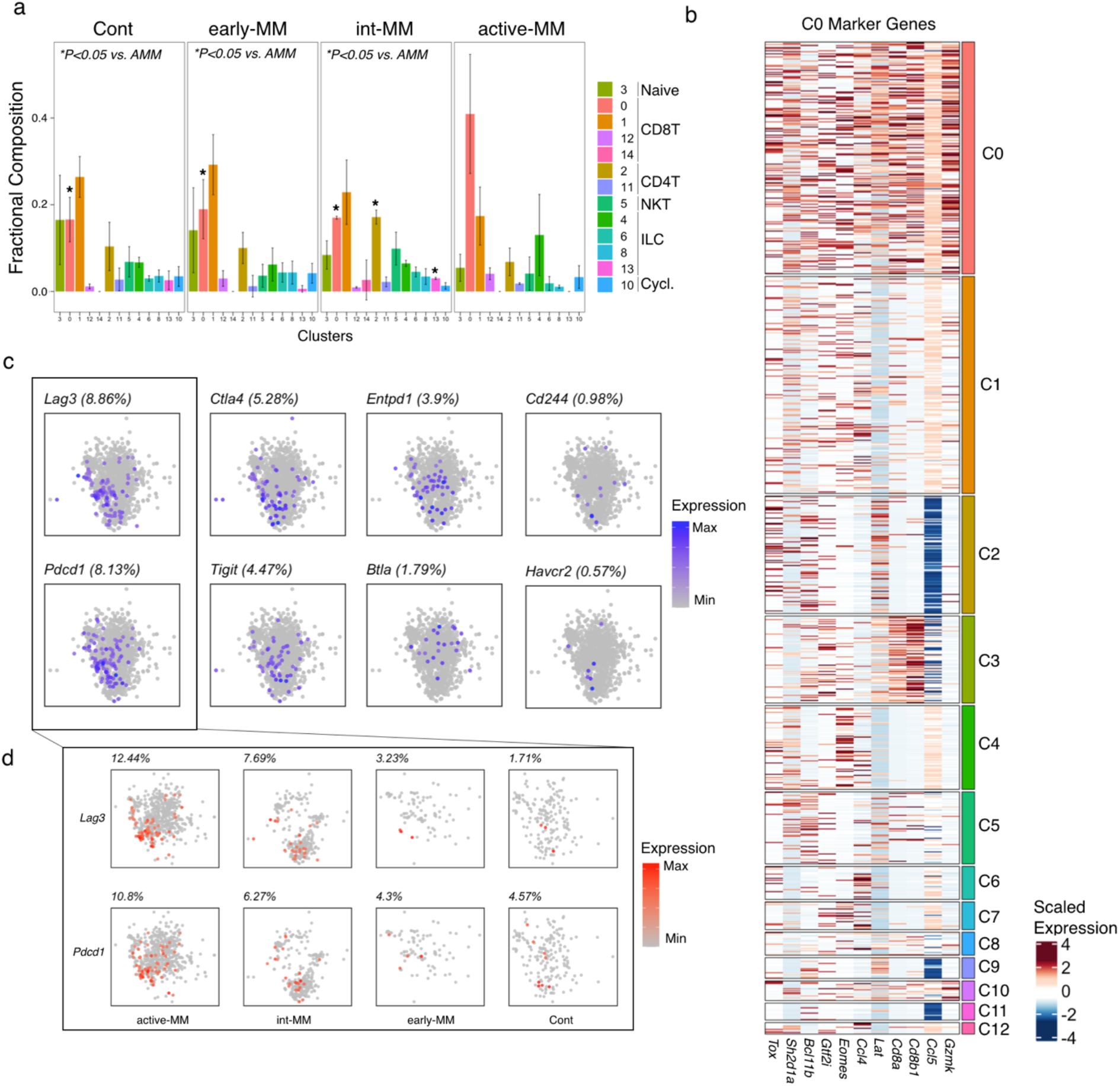
Defining T/NK subpopulations associated with defined disease stage groups. (a) Fractional composition of T/NK cell subpopulations, with data representing the average proportion within each disease stage cohort and error bars depicting standard deviation of the mean. Statistical comparisons were performed using a one-way ANOVA with Tukey multiple pairwise comparison and only P values for statistically significant comparisons are listed. (b) Heatmap depicting expression of C0 marker genes across all T/NK clusters (FDR<0.05). Data represent scaled expression values (any values outside a range of −2.5 to 2.5 were clipped). (c) UMAP visualization of log-normalized immune checkpoint receptor expression in C0 cells from all mice. (d) Disease-stage specific expression of Lag3 and Pdcd1 in C0 cells. Percent of cells expressing respective genes (UMI count ≥ 1) is listed above each plot in (c) and (d).

**Figure 10.**
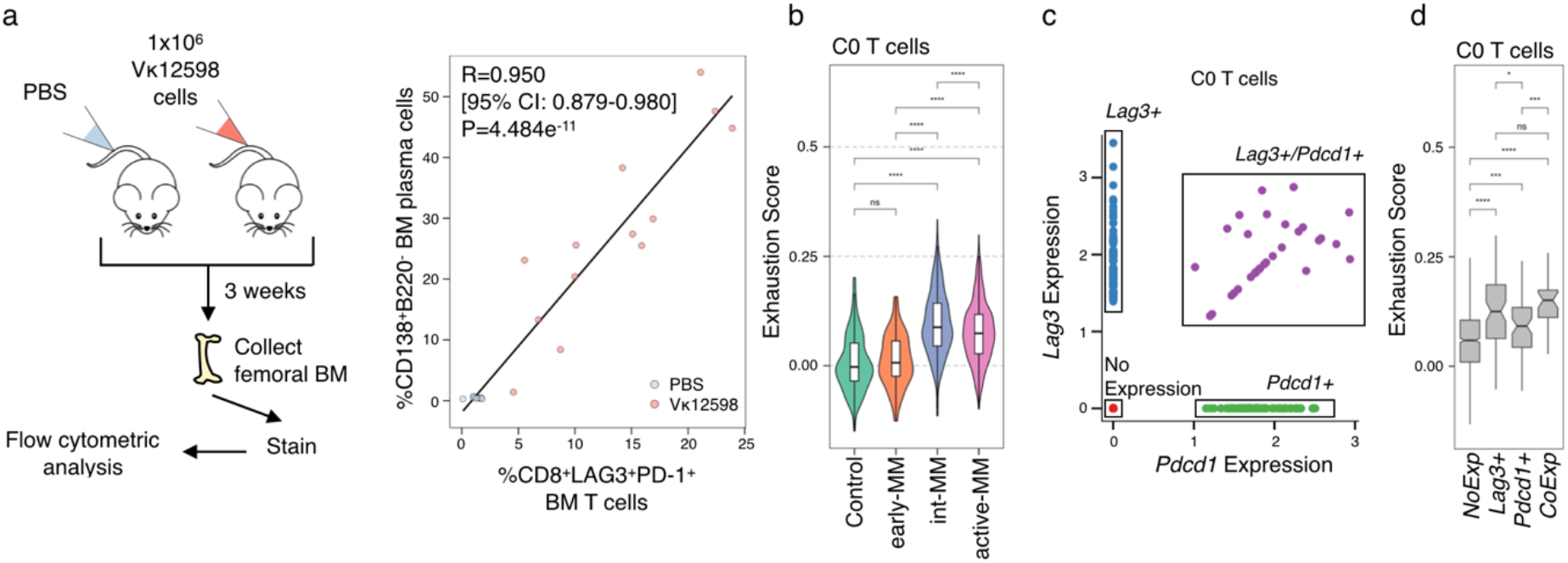
T cell exhaustion is detected early in Vκ*MYC disease progression and is associated with co-expression of immune checkpoint receptors. (a) Workflow for and results of validation experiments using the transplantable Vκ12598 model. Flow cytometric analyses of Vκ12598 samples, depicting correlation (Pearson’s R and regression line from linear model) between disease burden in BM (%CD138+B220−) and co-expression of PD-1 and LAG3 on CD8+ T cells. Each dot represents one mouse. (b) Distribution of T cell exhaustion gene signature scores in C0 cells across disease stage groups calculated using a gene set from Tirosh et al. [165]. (c) Scatter plot of *Lag3* and *Pdcd1* expression in C0 cells used to define expression groups for (d). (d) Distribution of Exhaustion gene signature scores in C0 cells amongst assigned expression groups based on *Lag3* and *Pdcd1* expression. Mean gene signature scores in (b) and (d) were compared across all groups using the Wilcoxon rank sum test (two-sided) corrected for multiple testing (Benjamini-Hochberg, *P<0.05, ***P<0.001, ****P<0.0001, ns=not significant).

### Combinatorial ICR blockade delays disease progression if administered early

Our data thus far supports that transcriptional shifts in the immune compartment occur early during the process of disease evolution and nominates potential drivers of progression. Therefore, we next aimed to evaluate whether early therapeutic intervention could delay disease progression focusing on our findings related to early onset of T cell exhaustion and co-expression of LAG3/PD-1. The rationale for this investigation is further supported by the observation that anti- (α)- PD-1 treatment in Vκ*MYC mice with overt disease fails to induce tumour regression [22]. We therefore treated Vκ12598-engrafted mice with combinatorial α-LAG3 and α-PD-1 monoclonal antibodies, initiating treatment 7 days post-tumour engraftment, before evidence of disease (Figure 11A). Disease burden was significantly reduced in mice treated with combinatorial α-LAG3/α-PD-1 compared to isotype-control treated mice as evidenced by lower serum M-protein after 3 weeks of treatment (Figure 11A-B, P=0.000094) and lower plasma cell infiltration in the BM (Figure 11C-D, P=0.00079) and spleen (Figure 11E-F, P=0.000096) at the time of sacrifice. Notably, disease burden was not significantly different between mice treated with either of the single-agent monoclonal antibodies and isotype control (Figure 11G). Finally, mice treated with combinatorial α-LAG3/α-PD-1 had significantly improved survival compared to mice treated with isotype controls (Figure 11H P=0.0013). These data therefore suggests that the associated increase in ICR-mediated T cell exhaustion observed by scRNA-seq reflects a causative mechanism of disease progression and supports dual blockade of these ICRs as a potential therapeutic strategy for high-risk precursor patients.

**Figure 11:**
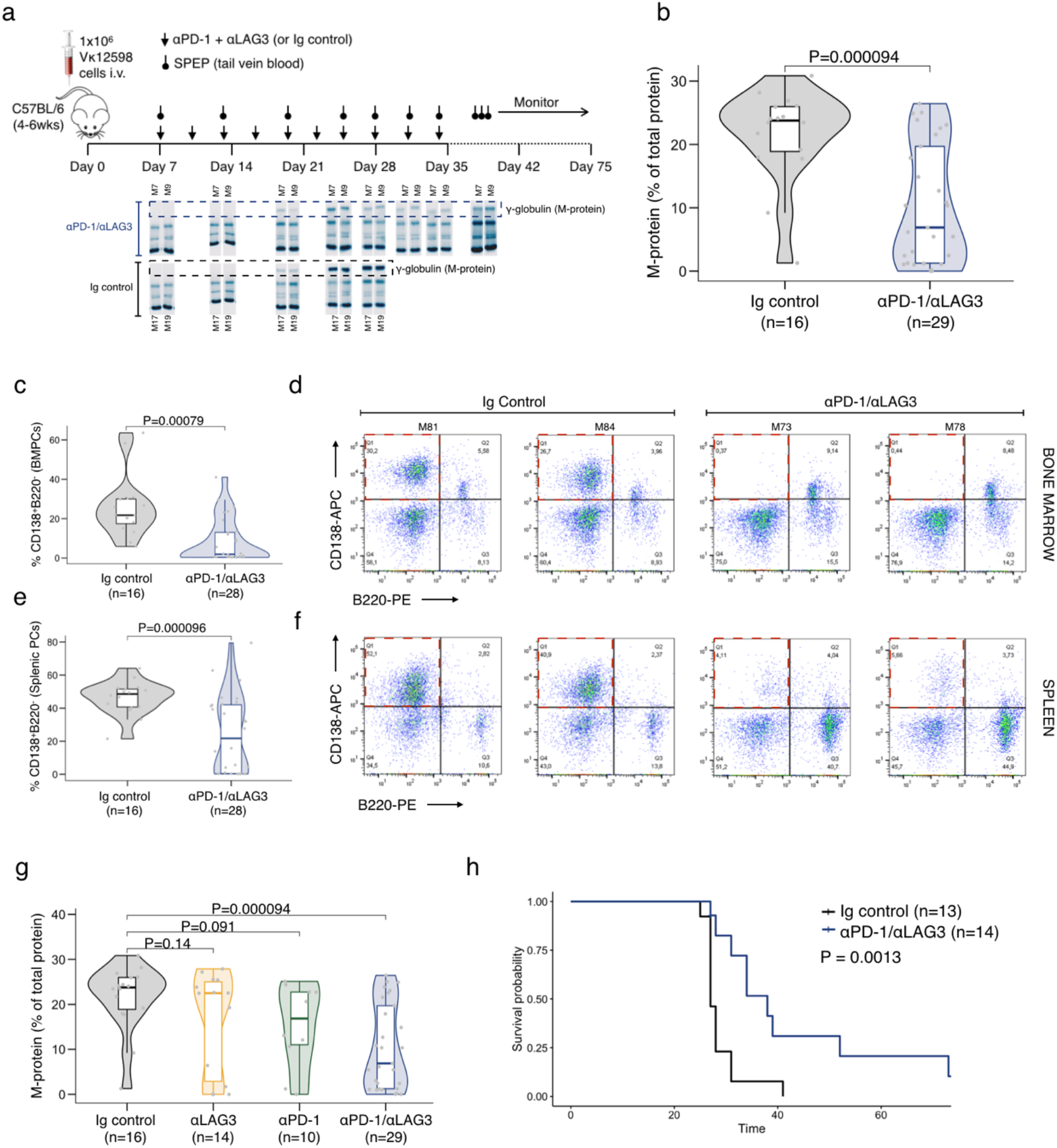
Early intervention with combinatorial ICR blockade can delay disease progression and improve survival. (a) Study design for in vivo combinatorial αLAG3/αPD-1 treatment experiments using Vκ12598 mice. (b) Serum M-protein measurements taken three weeks post-treatment initiation. (c) Bone marrow disease burden at time of sacrifice, as measured by flow cytometric analysis of %CD138+B220− plasma cells. (d) Scatter plots from flow cytometric analysis of BM disease burden (CD138+B220−) in representative mice treated with either Ig Control or αLAG3/αPD-1. Quadrants corresponding to myeloma/plasma cells are highlighted in red. (e) Splenic disease burden at time of sacrifice, as measured by flow cytometric analysis of %CD138+B220− plasma cells. (f) Scatter plots from flow cytometric analysis of splenic disease burden (CD138+B220−) in representative mice treated with either Ig Control or αLAG3/αPD-1. Quadrants corresponding to myeloma/plasma cells are highlighted in red. Statistical comparison of means in (b), (c), and (e) were performed using a two-sided t-test. (g) Serum M-protein measurements taken three weeks post-treatment initiation in mice treated with either Ig control, monotherapy αLAG3, monotherapy αPD-1 or combinatorial αLAG3/αPD-1 (Wilcoxon rank sum test (two-sided) corrected for multiple testing with Benjamini-Hochberg). (h) Kaplan-Meier survival plot of Vκ12598 mice treated with control or combinatorial αLAG3/αPD-1 (log-rank test).

## Discussion

Immune dysregulation plays a role in controlling disease progression in cancer [23], including myeloma [24–26]. However, it is only with the rapid uptake of single cell profiling that we are now beginning to unravel the diverse landscape of cell types, their molecular states, and dynamic interactions associated with disease processes in myeloma [14, 27–30]. Challenges towards these efforts remain, including extreme inter-patient variability and a paucity of samples from patients with early precursor disease. By employing a transgenic mouse model of myeloma, we performed a controlled, systematic analysis of immune dysregulation throughout progression. In doing so, we comprehensively characterize the features of the immune TME over the course of disease, from the earliest stages (prior to disease detection by traditional serological measurements), through intermediate and late-stage disease. This approach revealed novel immune-based targets for delaying disease progression, which we validated in vivo.

We acknowledge that the current study is not the first to employ scRNA-seq to profile immune alterations throughout disease progression [14, 28]. Using primary BM samples that span the myeloma disease spectrum, Zavidij et al. importantly reported that the immune TME is compromised early in MM [14]. Although our work did not identify all of the same changes in the immune TME as reported by Zavidij et al., our finding that T cell exhaustion occurs early supports their major conclusion that immune dysfunction is an early event in myeloma evolution. Our study further provides a unique perspective by surveying progression on a uniform genetic background.

Nonetheless, additional studies will be required to understand whether the compositional alterations and immune-associated changes throughout progression identified in our study are specific to a molecular subgroup or if they are an underlying feature of disease progression in all myeloma patients. To do this, future work will need to focus on applying single-cell approaches to larger patient cohorts.

Our analysis of the myeloid cell fraction identified several different populations that implicate IL-17 signaling in Vκ*MYC disease progression. Although we show elevated IL-17 signaling in Ly6c+ monocytes from the TME of mice with intermediate disease, we cannot say whether this merely reflects a bystander effect of IL-17 being present in the TME or if activation of IL-17 signaling in these cells has a direct causative role in progression. Nonetheless, previous work has established that inhibition of IL-17 in the TME delays disease progression and clearly supports IL-17 as an important player in the biology of precursor disease, both as a marker of high-risk disease and as a target for therapeutic intervention [11, 31]. In our study, IL-17 signaling was also activated in basophils and correlated with Il6 expression, a potent pro-MM cytokine [15, 16, 32]. Because basophils are typically depleted from myeloma samples during processing, little is known about their role in disease biology. However, it is well-known that basophils migrate to sites of inflammation, and these cells have been identified in human lung, gastric, pancreatic and ovarian cancer [33]. Thus, our findings of an IL-17-basophil-IL-6 axis activated during earlier stages of disease warrant further attention in future studies and provide rationale for investigating the role of cell types in disease progression that are traditionally disregarded during sample collection/processing.

In addition to providing insight into the molecular processes that underlie disease progression from early-MM to active disease, we aimed to nominate novel therapeutic interventions in the immune TME that could be exploited to delay disease progression. Towards this, we identified exhausted CD8+ T cells to be enriched in mice with intermediate and active disease and showed that these cells co-express the immune checkpoint receptors, LAG3 and PD-1. These cells also demonstrated upregulation of the transcription factor *Tox*, which has been shown to play a role in T cell exhaustion by positively regulating the expression of several ICRs [34]. We also demonstrated that co-inhibition of LAG3/PD-1 can delay disease progression and improve survival of mice if administered during early disease stages. These data are consistent with a model proposed by Jing et al., where reactivation of MM-specific T cells that were previously rendered dysfunctional by ICR proteins could be achieved by whole body irradiation-induced lymphopenia [35]. In this model, subsequent ICR blockade allowed reactivated T cells to remain functional and eliminate MM cells. Our study supports a similar model whereby ICR treatment allows reactive T cells to remain functional by administering early (versus by lymphodepletion). Thus, in certain biological contexts, there may be an advantage to using ICR blockade to prevent the onset of T cell dysfunction as opposed to reversing it once established.

Alternatively, the effectiveness of ICR blockade during earlier disease stages may be related to specific cell types that are present in precursor disease stages but subsequently lost during progression. For example, there is increasing support in the literature that a TCF1+ subset of exhausted T cells with “stem-like” features mediates the effects of α-PD-1 in models of chronic viral infection [36] and anti-tumour immunity [37]. Interestingly, a similar subset of TCF1hi memory T cells was recently described to be enriched in MGUS compared to MM [28]. Although we did not observe differential enrichment of a similar cell type in Vκ*MYC mice with early disease, this could be due to an insufficient number of cells profiled. Thus, future scRNA-seq studies could consider enriching samples for T cells specifically, as this would provide more insight into the heterogeneous transcriptional programs of dysfunctional T cells during precursor progression and preventative treatment. Moreover, given that PD-1 inhibitors have failed to demonstrate clinical activity as single agents in patients with active myeloma [38, 39], these findings have direct translation implications and support evaluating the clinical utility of this combination in pre-malignant or low tumour burden settings.

In conclusion, we present an in-depth analysis of immune dysregulation during disease evolution in the Vκ*MYC model, which provides insights into mechanisms driving progression to overt MM that may serve as biomarkers of high-risk smoldering myeloma. Our data also reveal tumour-extrinsic vulnerabilities that can be exploited as anti-MM therapies. We believe that findings from this study provide rationale for preventative treatment strategies in precursor disease stages of myeloma and more broadly, reveal critical insight into how immune cell populations contribute to disease evolution in cancer.

## Methods

### Mice and BM specimen processing

Animals used in this study were housed in pathogen-free facilities at either the Montreal University Health Centre or University Health Network under the following conditions: 21°C ambient temperature, 40-60% humidity, 12 hour dark/light cycle. All related experiments were approved by institutional Animal Care Committees and performed in accordance with the Canadian Council on Animal Care Guidelines (University Health Network Protocol #958.23, Montreal University Health Centre Protocol #2012-7242). For all scRNA-seq experiments, we employed the Vκ*MYC transgenic mouse model [12] cross-bred onto C57BL/KaLwRij given the latter model’s increased propensity for developing spontaneous monoclonal gammopathies and bone disease [40, 41]. To evaluate disease burden, we measured M-protein by SPEP. Mice were assigned to disease groups according to age and M-protein levels as follows (Figure 5A-B): early disease (early-MM: 27-33 weeks, not detected/trivial M-protein), intermediate-MM (int-MM, 49 weeks, trivial M-protein), and active-MM (61-74 weeks, major M-protein). Non-transgenic age-matched C57BL/KaLwRij mice (55-72 weeks) were also included to control for the effects of aging. For validation and in vivo treatment studies using the transplantable Vκ12598 model [42], we injected 1×10^6^ Vκ12598 cells in phosphate buffered saline (PBS), or PBS only, into 4-6 week old female C57BL/6J mice (tail vein). At the indicated time points, mice were sacrificed and BM material was extracted by flushing femurs and tibias with ice cold PBS. Cells were then dissociated by passing through a 23-gauge needle and debris removed using a 35 µm cell strainer. The resulting BM cells were subject to red blood cell lysis (ACK buffer), washed and resuspended in appropriate buffer for downstream analyses.

### Monitoring serum M-protein levels

To monitor the onset of disease in mice, 50 uL of tail vein blood was collected into microcuvettes (Sarstedt Inc, Newton, NC, USA) and centrifuged for 5 minutes at 10,000xg to separate serum. SPEP was performed with 0.5-1 μL of serum using the QuickGel System (Helena Laboratories, Beaumont, Texas, USA) according to manufacturer’s instructions. Densitometric quantification of bands was performed using Image J software (NIH, Open Source) as M-protein/total protein.

### Single-cell library preparation and sequencing

Single-cell suspensions were obtained from BM as described above and examined for cell number and viability using trypan blue and a Countess II automated counter (Thermo Fisher Scientific, Burlington, ON, Canada). Cell viability was greater than 70% for all samples described in this study. Single-cell libraries were constructed using the V2 chemistry kit from 10X Genomics (Pleasanton, CA, USA) according to manufacturer’s instructions. Libraries were sequenced on an Illumina HiSeq 2500 targeting 60,000 reads/cell at the Princess Margaret Genomics Centre (https://www.pmgenomics.ca).

### Sequence data pre-processing

The 10X Genomics CellRanger software suite (v2) was used to process raw sequencing reads, for alignment and to generate a digital gene expression (DGE) matrix of genes-by-cell counts. To account for the human MYC transgene in Vκ*MYC mice, the 10X Genomics GRCm38 genome reference package was supplemented with the GRCh38 MYC sequence and gene annotation. Total read counts for each cell barcode were calculated using BAMTagHistogram (Drop-seq Cookbook v1-2.12 [43]), followed by cell barcode calling using findInflectionPoint (dropbead v0.3 [44]). The resulting DGE matrices containing cell-associated barcodes only were merged for multi-sample analyses. Low-quality cells (<500 genes, <1000 transcripts (unique molecular identifier (UMI)), and/or >15% mitochondrial UMIs) and lowly expressed genes (expressed in less than 0.1% of the average number of cells per sample) were identified and removed from analysis. Finally, suspected doublets were removed from the dataset if identified using doubletFinder [45] (v2.0.3).

### Single-cell RNA-seq secondary analyses

All subsequent analysis steps were performed using Seurat [46] (v3.2.1) in R v3.6.1 unless otherwise stated. This included log-normalization (NormalizeData), identification of top 3000 variably expressed genes (FindVariableFeatures, “vst” method) and data scaling (ScaleData, regression on the proportion of mitochondrial transcripts). The top variably expressed genes were used for principal component analysis (PCA, RunPCA), with significant principal components identified using KneeArrower (v0.1.0). Data were embedded in two-dimensional space using t-distributed stochastic neighbor embedding (t-SNE) [47] and/or Uniform Manifold Approximation and Projection (UMAP) [48]. Batch effect correction was applied using Harmony [49] (group.by.vars = sequencing batch, theta = 0.5) prior to clustering with the graph-based algorithm implemented by Seurat.

Cell types were assigned to each cluster using SingleR (v1.0.6) [13], which automatically annotates scRNA-seq expression data using a reference data set with known labels. Since our data set was derived from mouse BM cells, we used the ImmGen [17] reference data set (ref=immgen), which is comprised of bulk gene expression profiles from highly purified mouse immune and hematopoietic cell populations (GSE15907, GSE37448). Clusters were assigned to a “fine.label” cell type if it represented the majority proportion, and then adjustments were made based on visual inspection of the “main.label” cell type.

Differential gene expression was assessed with FindMarkers or FindAllMarkers using the Wilcoxon rank sum test and a significance of P<0.05 after Bonferroni correction. Enrichment analysis was performed using Enrichr [50, 51]. Gene signature scores were computed with AddModuleScore.

### Flow cytometry

Samples of 0.5-1 × 106 cells were suspended in staining buffer (PBS, 2% FBS, 0.5mM EDTA) containing the indicated fluorophor-conjugated antibodies (CD138-APC [281-2] and B220-PE [RA3-6B2] purchased from BD Biosciences; CD3-FITC [17A2], CD8-APC-CY7 [53-6.7], PD-1-APC [RMPI-30] and LAG3-PE [C9B7W] purchased from Biolegend). After staining for 30 minutes at room temperature, cells were washed and re-suspended in staining buffer containing viability dye (4′,6-diamidino-2-phenylindole (DAPI) or 7-Aminoactinomycin D (7AAD)). Data were acquired on a Canto II cytometer (BD Biosciences, San Diego, CA, USA) and analyzed using Flowjo (v10.7.1).

### In vivo antibody treatment

Anti-mouse LAG3, PD-1 and isotypic IgG1-control monoclonal antibodies (provided generously by Bristol Myers Squibb) were administered via intraperitoneal injection in doses of 10 mg/kg each for LAG3/PD-1 and 20 mg/kg for IgG1 isotype control. For monotherapy, 10 mg/kg of single-agent monoclonal antibodies were combined with 10 mg/kg IgG1. These were administered every 3 days beginning seven days post-engraftment of Vκ12598 cells for a total of 10 doses. Disease was monitored by SPEP and mice were sacrificed when M-protein represented >20% of total serum proteins or physical signs of disease were observed. Upon sacrifice, BM was extracted as described above.

### Statistics

Statistical tests performed are as indicated in figure legends. Sample size was determined by the availability of subjects for scRNA-seq, but a minimum of 3 samples for each disease stage group was decided upon upfront to capture potential biological heterogeneity between tumours. Measurements were taken from distinct samples unless otherwise stated. Boxplots represent the distribution of each measurement within defined groups, where the central rectangle spans the interquartile range, the central line represents the median, and “whiskers” above and below the box show the value 1.5x the interquartile range.

## Data and code availability

All data described in this study were deposited to the National Center for Biotechnology Information Gene Expression Omnibus (GSE134370) and Sequence Read Archive (SRP214856). Data generated from this study is also available through the interactive single-cell portal CReSCENT (CRES-P32, https://crescent.cloud/) [52]. Code supporting this study is available at github.com/pughlab/scMM_VkMYC_2020_bin.

